# Radiosynthesis automation, non-human primate biodistribution and dosimetry of K^+^ channel tracer [^11^C]3MeO4AP

**DOI:** 10.1101/2023.03.28.534386

**Authors:** Yu-Peng Zhou, Moses Q. Wilks, Maeva Dhaynaut, Nicolas J. Guehl, Sung-Hyun Moon, Georges El Fakhri, Marc D. Normandin, Pedro Brugarolas

## Abstract

**Purpose:** 4-Aminopyridine (4AP) is a medication for the symptomatic treatment of multiple sclerosis. Several 4AP-based PET tracers have been developed for imaging demyelination. In preclinical studies, [^11^C]3MeO4AP has shown promise due to its high brain permeability, high metabolic stability, high plasma availability, and high *in vivo* binding affinity. To prepare for the translation to human studies, we developed a cGMP-compliant automated radiosynthesis protocol and evaluated the whole-body biodistribution and radiation dosimetry of [^11^C]3MeO4AP in non-human primates (NHPs).

**Methods:** Automated radiosynthesis was carried out using a GE TRACERlab FX-C Pro synthesis module. One male and one female adult rhesus macaques were used in the study. A high-resolution CT from cranial vertex to knee was acquired. PET data were collected using a dynamic acquisition protocol with 4 bed positions and 13 passes over a total scan time of ∼150 minutes. Based on the CT and PET images, volumes of interest (VOIs) were manually drawn for selected organs. Non-decay corrected time-activity curves (TACs) were extracted for each VOI. Radiation dosimetry and effective dose were calculated from the integrated TACs using OLINDA software.

**Results:** Fully automated radiosynthesis of [^11^C]3MeO4AP was achieved with 7.3 ± 1.2 % (n = 4) of non-decay corrected radiochemical yield within 38 min of synthesis and purification time. [^11^C]3MeO4AP distributed quickly throughout the body and into the brain. The organs with highest dose were the kidneys. The average effective dose of [^11^C]3MeO4AP was 4.27 ± 0.57 μSv/MBq. No significant changes in vital signs were observed during the scan.

**Conclusion:** The cGMP compliant automated radiosynthesis of [^11^C]3MeO4AP was developed. The whole-body biodistribution and radiation dosimetry of [^11^C]3MeO4AP was successfully evaluated in NHPs. [^11^C]3MeO4AP shows lower average effective dose than [^18^F]3F4AP and similar average effective dose as other carbon-11 tracers.

## INTRODUCTION

4-Aminopyridine (4AP) is a voltage-gated potassium channel blocker (**Figure 1**), which binds inside the pore of potassium channel under protonated condition [1, 2]. 4AP has been approved by the U.S. Food and Drug Administration for the symptomatic treatment of multiple sclerosis (MS) [3-5]. Upon demyelination, axonal potassium channels K_v_1.1 and K_v_1.2 normally located under the myelin sheath become exposed and increase in expression. 4AP binds to the potassium channels in demyelinated axons, reducing the abnormal efflux of K^+^ ions and restoring axonal conduction [6, 7]. Several 4AP based PET tracers have been developed by our group (**Figure 1**) [8-11]. The fluorine-18 based PET tracer [^18^F]3-fluoro-4-aminopyridine ([^18^F]3F4AP) has been characterized in healthy non-human primates and healthy human subjects, showing selective binding to potassium channels, high brain penetration, high metabolic stability, high plasma availability, high reproducibility, high specificity, and fast kinetics [8]. In addition, [^18^F]3F4AP showed high sensitivity to a traumatic brain injury (TBI) in a non-human primates [12]. Clinical trials of [^18^F]3F4AP in people with multiple sclerosis (NCT04699747), neurodegeneration and traumatic brain injury patients (NCT04710550) are currently underway.

**Figure 1.**
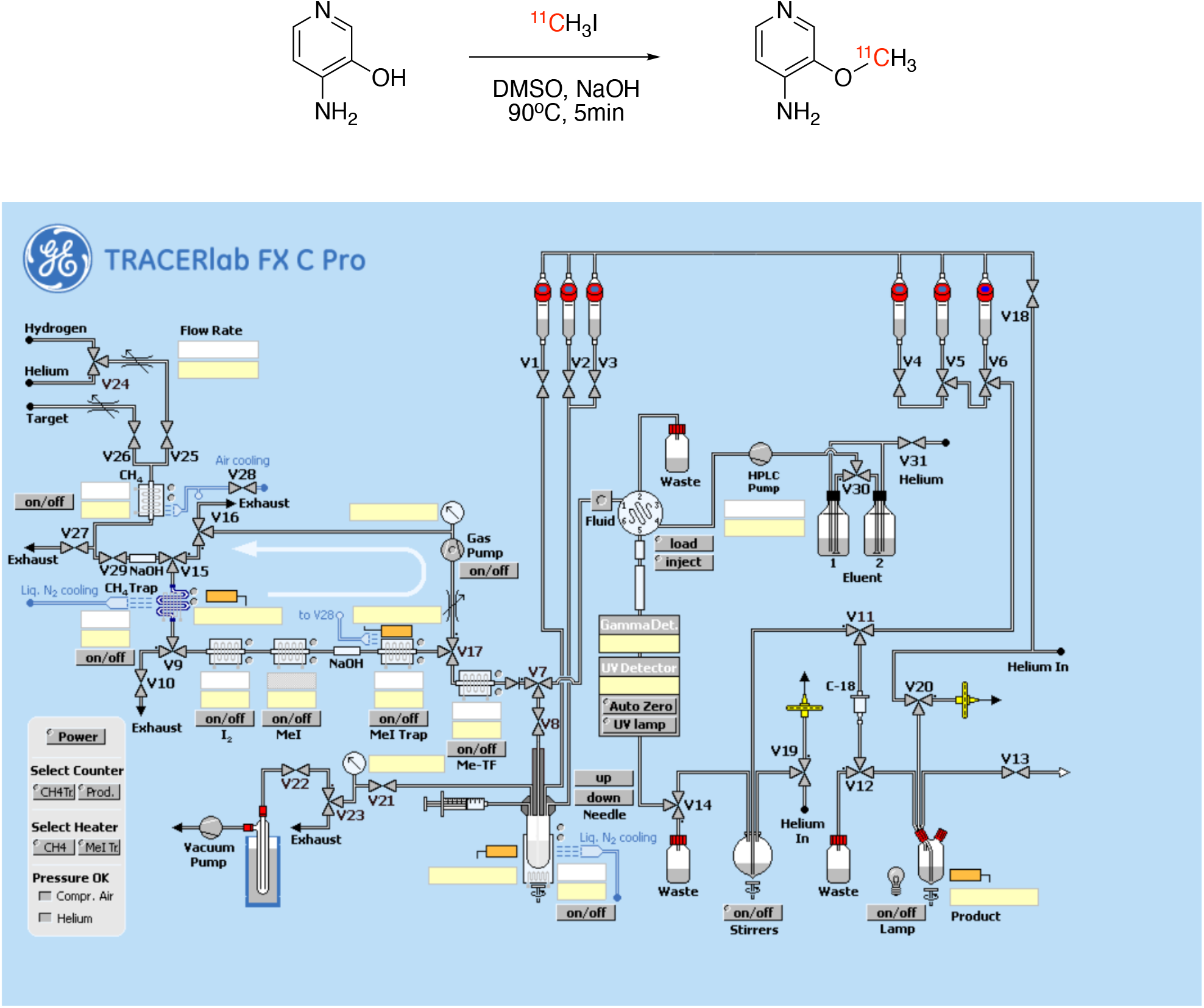
Radiosynthesis of [^11^C]3MeO4AP (top), and control panel of GE TRACERlabT FX C Pro synthesizer (bottom).

Carbon-11 is one of the most commonly used PET radionuclides. The half-life of carbon-11 (t_1/2_ = 20.3 min) is shorter than that of fluorine-18 (t_1/2_ = 110 min). Due to their shorter half-life, carbon-11 PET tracers generally display lower effective doses and offer the opportunity to scan the same subject twice on the same day [13-18]. Several carbon-11 labeled 4AP-based PET tracers have been reported including 3-[^11^C]trifluoromethyl-4-aminopyridine ([^11^C]3CF_3_4AP) [9, 19], 3-[^11^C]methoxy-4-aminopyridine ([^11^C]3MeO4AP) [10], and 3-[^11^C]methyl-4-aminopyridine ([^11^C]3Me4AP) [11] (**Figure 1**). Among them, [^11^C]3MeO4AP was the only one that showed marked uptake in a three-year old focal traumatic brain injury in a rhesus macaque. Additionally, [^11^C]3MeO4AP showed several promising properties such as high brain permeability, high metabolically stability, high plasma availability, and higher *in vivo* binding affinity and specificity to potassium channels compared with [^18^F]3F4AP [10]. The properties mentioned above make [^11^C]3MeO4AP a promising candidate for imaging demyelinating diseases. In order to bring [^11^C]3MeO4AP to a clinical study, the cGMP (Current Good Manufacturing Practices) production of PET tracer and the radiation dosimetry evaluation are required to fulfill the IND (Investigational New Drug) application requirement. In this paper, we describe the cGMP compliant automated radiosynthesis and quality control of [^11^C]3MeO4AP and its biodistribution and radiation dosimetry in non-human primates.

**Scheme 1.**
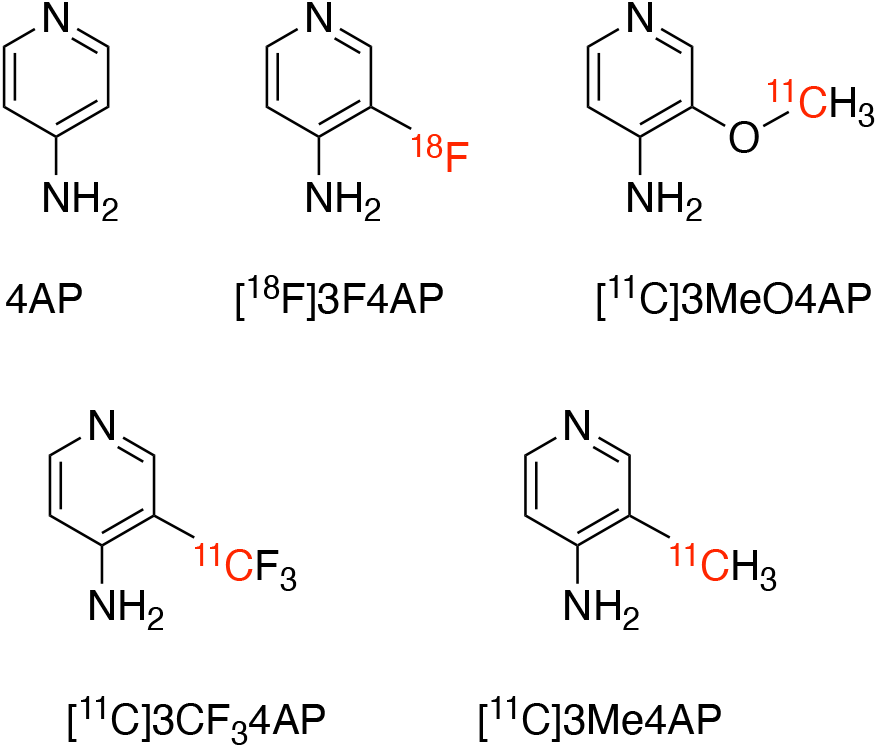
Chemical structures of 4-aminopyridine (4AP) and related PET tracers.

## EXPERIMENTAL SECTION

### Compliance

All experiments involving non-human primates were performed in accordance with the U.S. Department of Agriculture (USDA) Animal Welfare Act and Animal Welfare Regulations (Animal Care Blue Book), Code of Federal Regulations (CFR), Title 9, Chapter 1, Subchapter A, Part 2, Subpart C, §2.31. 2017. Animal experiments were approved by the Animal Care and Use Committee (IACUC) at the Massachusetts General Hospital (MGH).

### Non-human primates

One male adult (M1) and one female adult (M2) rhesus macaque were used in this study. Animal body weights on the days of imaging were 13.68 kg (Male) and 9.74 kg (Female). Prior to each imaging session, animals were sedated with ketamine/xylazine (10/0.5 mg/kg IM) and were intubated for maintenance anesthesia with isoflurane (1-2% in 100% O_2_). A venous catheter was placed in the saphenous vein for radiotracer injection and, an arterial catheter was placed in the posterior tibial artery for blood sampling. The animal was positioned on a heating pad on the bed of the scanner for the duration of the study. During 150 min of PET scan, vital signs including temperature, blood pressure and oxygen saturation, heart rate, respiratory rate, and exhaled CO_2_ were monitored.

### PET tracer production

#### Automated radiosynthesis of [^11^C]CH_3_I

The proton bombardment (40 μA, 3-7 min) of nitrogen gas in the presence of oxygen (1%) generated [^11^C]CO_2_ via ^14^N(p,α)^11^C nuclear reaction. The cyclotron generated [^11^C]CO_2_ was transferred into a GE TRACERlab FX MeI module or TRACERlab FX C Pro synthesis module and then mixed with hydrogen. The mixture was passed over nickel at 350 °C to produce [^11^C]CH_4_, which was then circulated and passed through a high-temperature iodine bed. The sublimated iodine was reacted with [^11^C]CH_4_ in gas phase to generate [^11^C]CH_3_I, which was first trapped in MeI trap and then released, transferred into a V-vial (semi-automated method) or to the reactor (fully automated method using TRACERlab FX C Pro synthesis module) for the radiomethylation reactions.

#### Semi-automated radiosynthesis of [^11^C]3MeO4AP

3-5 mg of 3-hydroxyl-4-aminopyridine, 300 μL DMSO, and 5 μL of 5N NaOH solution were added into a 5 mL V-vial. The mixture was vortexed for 1 minute and nitrogen gas was bubbled through the solution for 3 minutes until the solution turned pink. The produced [^11^C]CH_3_I was then bubbled into the mixture for 3 minutes at room temperature. All the needles were removed and the sealed V-vial was heated at 90 °C for 5 minutes. When reaction was completed, 2.5 mL of H_2_O was added via syringe and the mixture was transferred to the prep-HPLC for purification. The tracer was purified using a semiprep HPLC column (Waters XBridge C18 10 × 250 mm) using 10 mM sodium phosphate (pH 8) mobile phase containing 5% ethanol. The HPLC fraction containing the product (approx. 6-8 min) was diluted with 10 mL of 0.9% sodium chloride for injection. The identity and purity of [^11^C]3MeO4AP was confirmed by analytical HPLC with co-injected nonradioactive reference as standard.

#### Fully automated radiosynthesis of [^11^C]3MeO4AP

The fully automated radiosynthesis was carried out using a GE TRACERlab FX C Pro synthesis module. DMSO solution of 3-hydroxyl-4-aminopyridine containing base was prepared similar to the semi-automated radiosynthesis method. The mixture was preloaded in the reactor of the synthesis module. 1 mL of H_2_O was loaded into vial-3 (Figure 1). The produced [^11^C]CH_3_I was send to reactor via valve-8 at room temperature. The mixture was heated at 90 °C for 5 minutes. 1 mL H_2_O from vial-3 was added to the mixture and the solution was transferred to the prep-HPLC for purification using the same condition. The whole process was pre-programmed and run automatically. The identity and purity of [^11^C]3MeO4AP was confirmed by the same method used in semi-automated radiosynthesis protocol.

#### Quality control (QC) tests

the following QC tests were carried out as follows: Appearance was checked by visual inspection. Radiochemical identity was confirmed by coelution of nonradioactive 3MeO4AP and [^11^C]3MeO4AP on radioHPLC. Molar activity was calculated by dividing the dose measured in a dose calibrator by the mass calculated using a calibration curve from HPLC. Radiochemical purity was calculated as the ratio of the area under the curve (AUC) of the product peak on radioHPLC to all other radioactive peaks. Radionuclidic identity was confirmed by calculating the half-life from two radioactivity measurements >10 min apart using a dose calibrator. Chemical purity was estimated by integrating all the UV active peaks on HPLC at 254 nm. Finally, the pH was measured using pH test strips.

### PET tracer administration

Radiotracer solution (10 mL) was administered via the lateral saphenous vein over a 3-minute infusion. All injections were performed using syringe pumps (Medfusion 3500). After administration of the dose, the catheter was flushed with 10mL of saline and the residual activity in the syringe and catheter measured to calculate the injected dose.

### Image acquisition protocol

Imaging was performed on a GE Discovery MI PET/CT scanner. Subjects were positioned on the scanner bed and a CT from the cranial vertex to the knee was acquired. Based on the CT images, 4 bed positions with overlapping edges were selected for PET acquisition (25 cm per bed position with 2.5-5 cm overlap on each end). PET images were acquired over a period of 2.5 hours. The PET acquisition protocol consisted of a series of static images at each bed position of increasing duration starting upon administration of the tracer. The full acquisition protocol was as follows: high resolution CT, 5 passes × 1 min/bed, 5 passes × 3 min/bed, 3 passes × 5 min/bed. After completion of the scan, the PET data was reconstructed using the scanner’s OSEM with PSF and TOF modeling reconstruction algorithm with 34 subsets and 2 iterations applying the corrections for scatter, attenuation, deadtime, random coincidence and scanner normalization.

### Blood sampling

1 mL venous blood samples were taken at approximately 3, 5, 10, 15, 30, 60, 90, 120 and 150 min post-injection. Blood samples’ radioactivity was counted using a calibrated gamma counter.

### Image analysis and dosimetry calculation

Based on the high resolution CT and PET images, volumes of interest (VOIs) were manually drawn for the following tissues: adrenals, brain, breasts, gall bladder, small intestine, upper and lower large intestine, stomach, heart contents, heart muscle, kidney, liver, lung, muscle, ovaries, pancreas, red marrow, trabecular and cortical bone, spleen, testes, thymus, thyroid, urinary bladder and uterus. Time-activity curves (TACs) were extracted for each VOI. For dosimetry calculation, TACs were uncorrected for decay and extrapolated to ten half-lives after injection by assuming that any further decline in radioactivity occurred only due to physical decay with no biological clearance. Radiation dosimetry and effective dose were calculated from the integrated TACs using OLINDA software.

## RESULTS

### Automation of radiosynthesis

The semi-automated radiosynthesis of PET tracer [^11^C]3MeO4AP was achieved using a similar method as previously reported [10]. Starting from 10.73 GBq – 16.65 GBq (290 mCi – 450 mCi) of [^11^C] CH_3_I, 1.37 GBq – 1.89 GBq (37 mCi – 51 mCi) of [^11^C]3MeO4AP was synthesized in 11.3 ± 2.1 % (n = 4) non-decay corrected radiochemical yield and >99% of radiochemical purity in ∼45 min of synthesis and purification time.

In order to fulfill the cGMP requirements, a fully automated radiosynthesis method was developed using GE TRACERlab FX C Pro automatic synthesizer (**Figure 1**). The module contains a [^11^C]CH_4_ synthesis module, a needle reactor, a prep-HPLC, and a formulation system. The radiosynthesis of [^11^C]3MeO4AP was preprogrammed and ran automatically. A typical automated radiosynthesis starts from 5.55 GBq – 6.66 GBq (150 mCi – 180 mCi) of [^11^C]CH_3_I and yields 0.38 GBq – 0.46 GBq (10.4 mCi – 12.47 mCi) of [^11^C]3MeO4AP in 7.3 ± 1.2 % (n = 4) of non-decay corrected radiochemical yield and 99% of radiochemical purity in 38 min of synthesis and purification time. **Table 1** summarizes the QC tests carried out and their results.

**Table 1.** Quality control (QC) results of [^11^C]3MeO4AP productions (n = 4).

| Test | Result | Acceptance criteria | Method |
| --- | --- | --- | --- |
| Appearance | Pass | Clear and Colorless solution | Visual inspection |
| Radiochemical identity | $3.3 \pm 0.1$ min | The retention time within 0.2 min of standard | Coinjection of non-radioactive 3MeO4AP and [$^{11}\text{C}$]3MeO4AP using radioHPLC |
| Molar activity | $51.8 \pm 3.7$ GBq/$\mu\text{mol}$<br>($1.4 \pm 0.1$ Ci/$\mu\text{mol}$) | $> 37$ GBq/$\mu\text{mol}$<br>(1 Ci/$\mu\text{mol}$) | Dose calibrator and HPLC |
| Radiochemical purity | $99 \pm 1$ % | $> 95\%$ | RadioHPLC |
| Radionuclidic identity | Pass | Half-life = $20.3 \pm 1$ min | Dose calibrator |
| Chemical purity | Pass | $< 50.0$ mAU per 10 $\mu\text{L}$ of final dose | Analytical HPLC injection 10 $\mu\text{L}$ of final dose |
| pH | $7 \pm 0.5$ | 5-8 | pH test strips |

### Biodistribution and radiation dosimetry of [^11^C]3MeO4AP

Two monkeys (one male, one female) were used in the study in order to be able to assess the dosimetry to reproductive organs. The subject characteristics are shown in table 2.

**Table 2.** Summary of subject information.

| Rhesus macaques | Sex | Weight | Height | Injected activity | Effective Dose |
| --- | --- | --- | --- | --- | --- |
| M1 | Male | 13.68 kg | 1.2 m | 293.4 MBq (7.93mCi) | 1.26 mSv |
| M2 | Female | 9.74 kg | 0.91 m | 290.5 MBq (7.85mCi) | 1.24 mSv |

As it can be seen from figure 2 (maximum intensity projections) and figure 3 (whole body sagittal slices) [^11^C]3MeO4AP quickly distributed widely throughout the body. Accumulation of tracer is clearly visible in the urinary bladder, kidneys, liver, thyroid, brain, salivary glands, and spinal vertebras. Figure 4 shows the TACs of blood obtained by gamma counting of serial arterial blood samples as well as a VOI placed in the heart left ventricular chamber (Fig 4A). In addition, TACs were extracted from VOIs placed in brain, thyroid, muscle, and bone (Fig. 4B); liver, stomach, spleen, and bone marrow (Fig. 4C) and kidneys and bladder (Fig. 4D).

**Figure 2.**
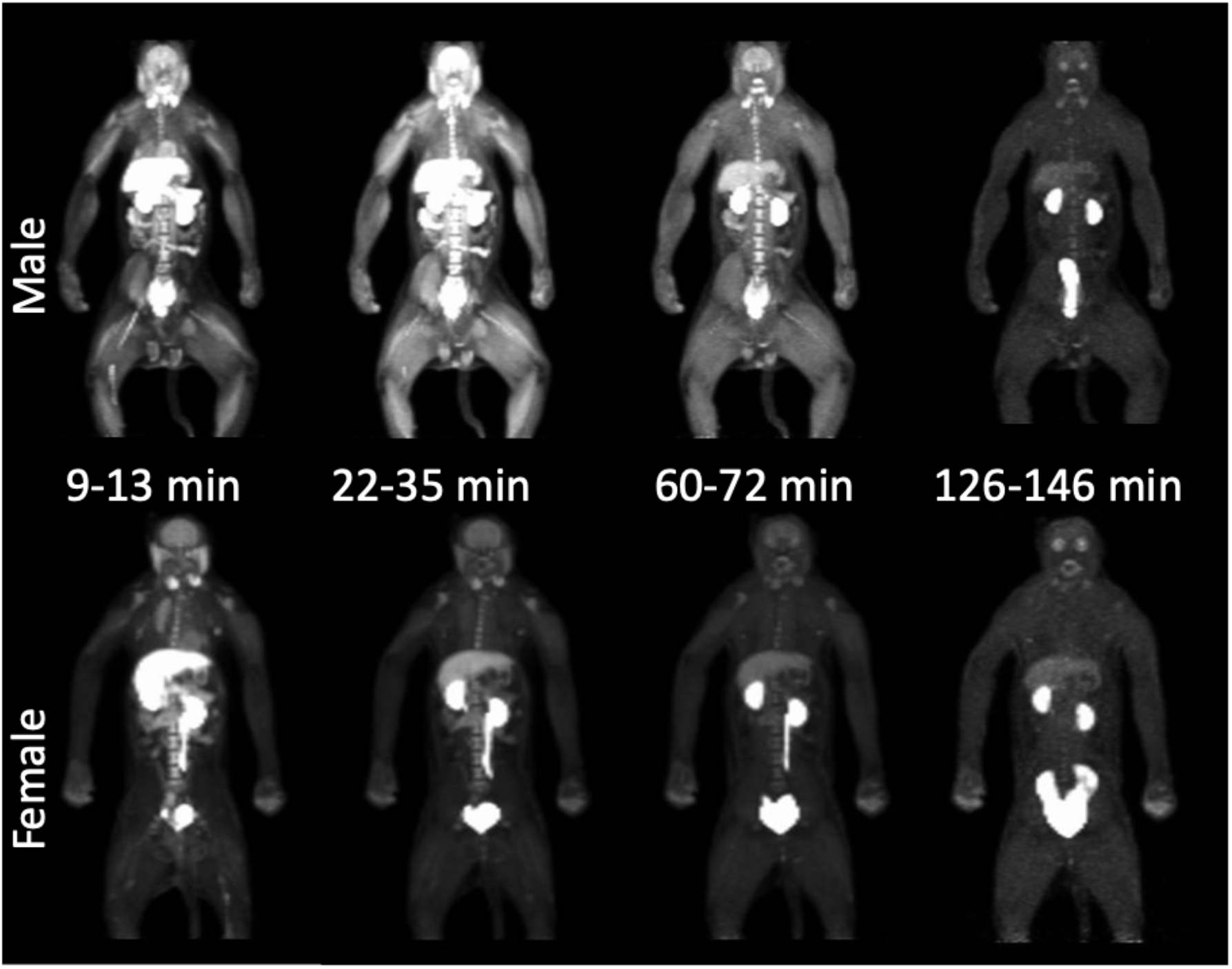
Whole-body maximum intensity projection (MIP) PET images (9-13 min, 22-35 min, 60-72 min, and 126-146 min) of rhesus macaques (male: top, female: bottom).

**Figure 3.**
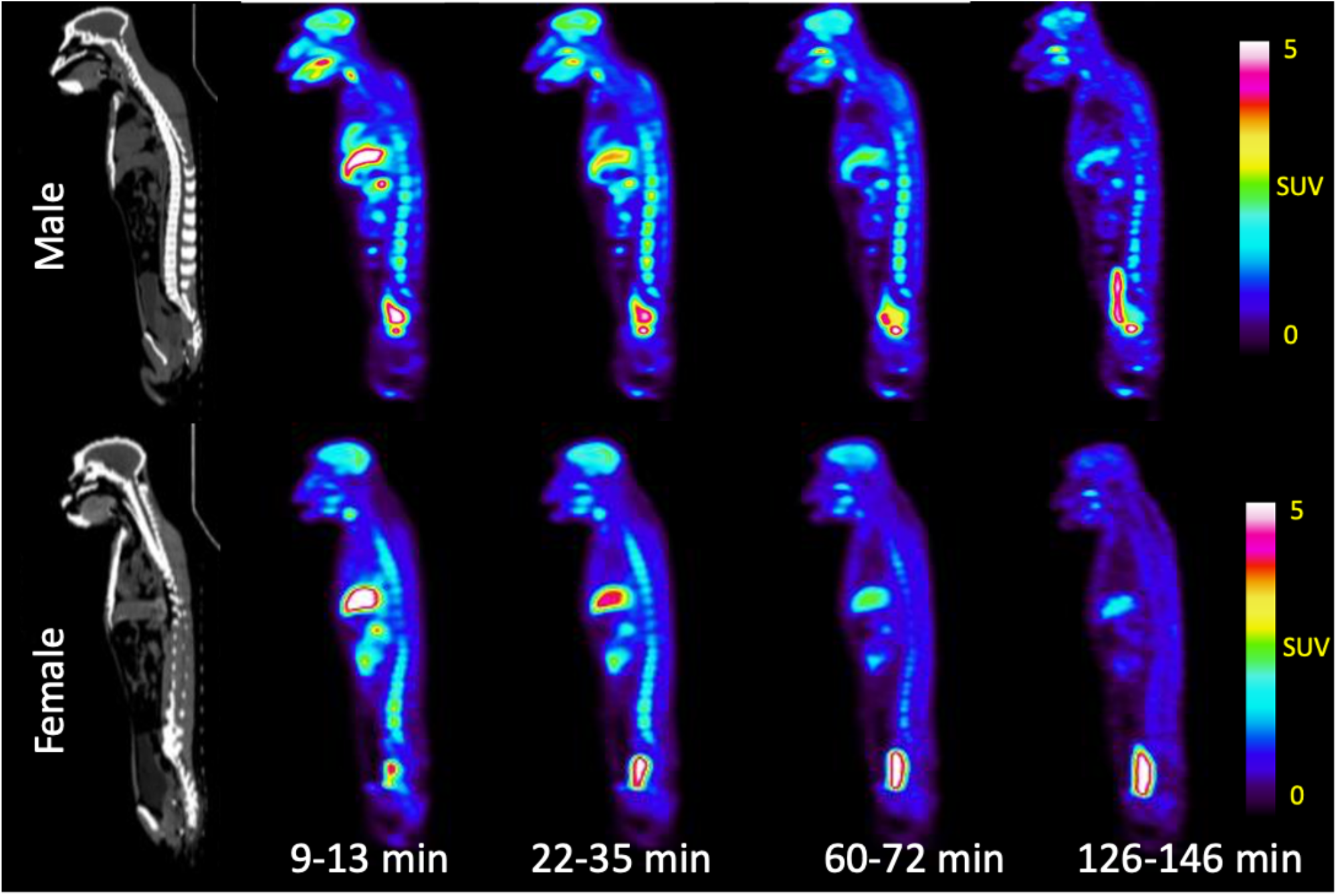
Representative whole-body CT and PET images (9-13 min, 22-35 min, 60-72 min, and 126-146 min) of rhesus macaques (Sagittal view, male at top, female at bottom)

**Figure 4.**
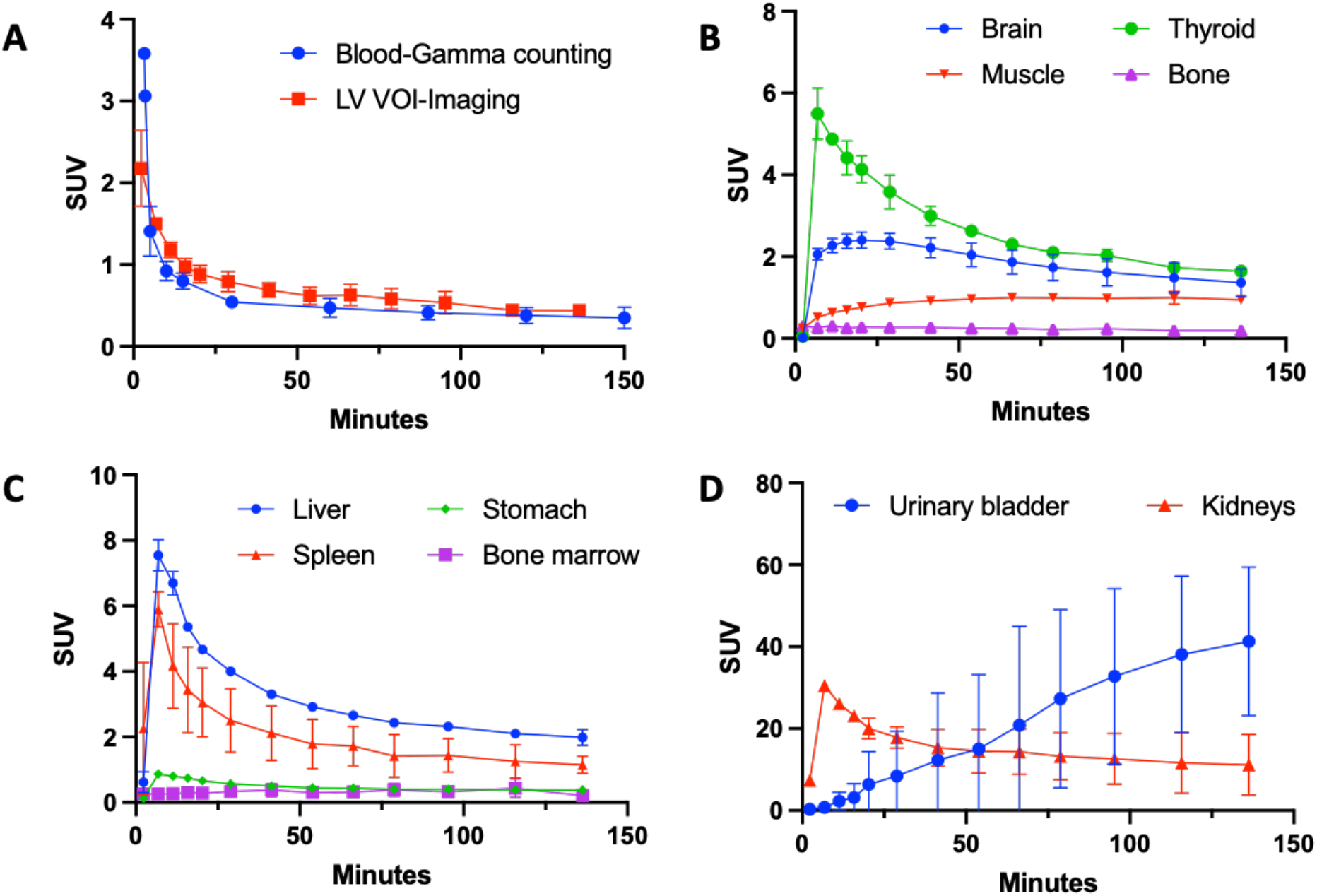
**A**: Whole blood decay-corrected time-activity curves measured by venous blooding samples gamma counting and left ventricle (LV) VOI of PET imaging. **B-D**: Decay corrected time-activity curves of selected organs.

The concentration in blood peaked at 0-3 min post injection and then it quickly decreased. The blood radioactivity clearance was fitted using a two-phase decay model indicating fast washout (male: t_½ fast_ = 0.68 min, t_½ slow_ = 9.3 min; female: t_½ fast_ = 0.50 min, t_½ slow_ = 19.1 min). Comparison of the blood measured by gamma counting and from a VOI placed in the hearth left ventricle showed very similar results highlighting the accuracy of both methods. From the images and TACs, it appears that the tracer is primarily cleared through the kidneys with the signal in the urinary bladder surpassing that of the kidneys after ∼ 55 min (Fig. 4B). In most organs (e.g. kidneys, liver, spleen, stomach, thyroid, etc.) maximum SUV was reached during at 2-11 min post-injection. Meanwhile, the SUV of muscle increases until 20 min post injection and remains stable at SUV ≈ 1 until the end of imaging.

Consistent with previous results [10], the pharmacokinetics in the brain were slower than other organs, reaching a whole brain SUV ≈ 2.4 at 16-29 min post-injection followed by a slow decrease to SUV ≈ 1.4 by the end of imaging (150 min post injection). The brain kinetics can also be observed in horizontal brain slices (Fig. 5), which show higher signal in grey matter than white matter as previously described [10]. There were no significant differences observed between male and female.

**Figure 5.**
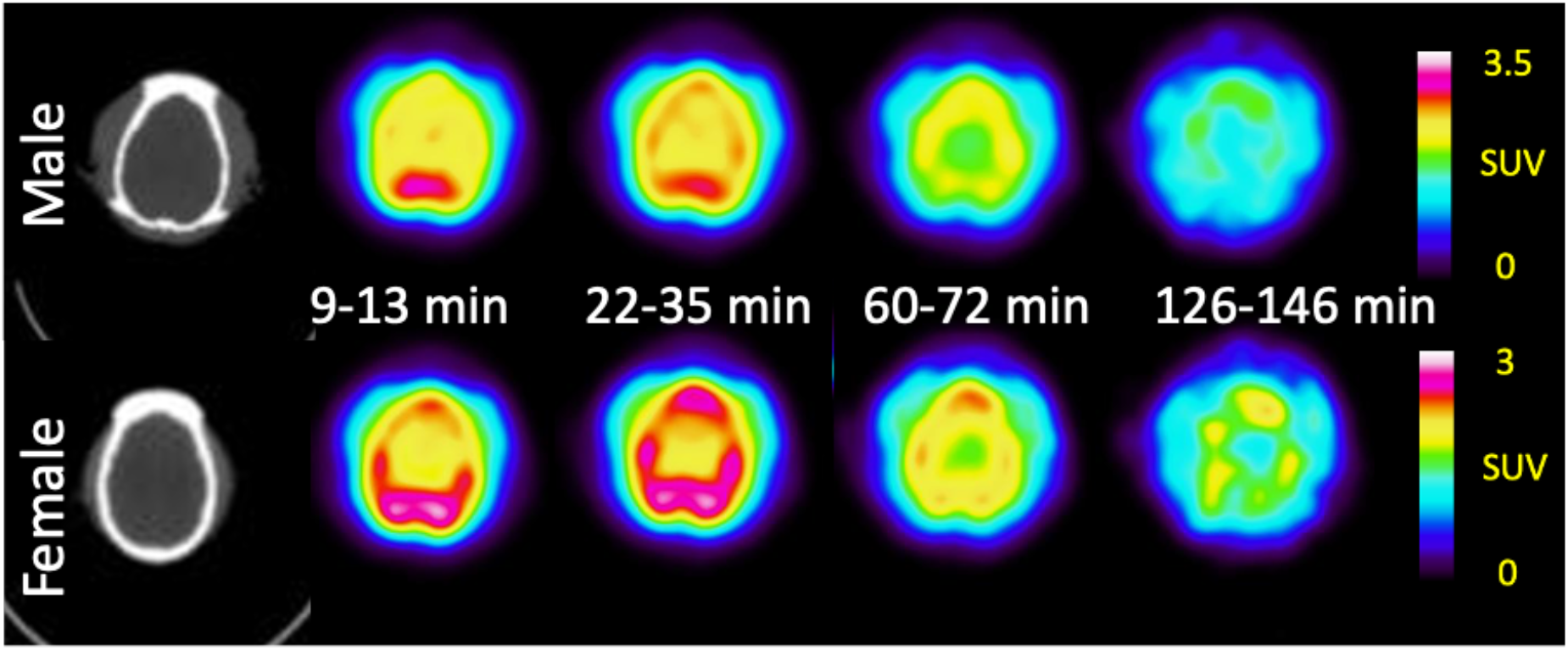
Representative brain CT and PET images (9-13 min, 22-35 min, 60-72 min, and 126-146 min) of rhesus macaques (male-top, female-bottom).

From the integration of the non-decay corrected TACs, the residence time of selected organs were calculated (Table 3). The brain, heart content (blood), kidneys, liver, and muscle of both male and female animal show relative longer residence time ranging from ∼0.02 MBq*h/MBq to ∼0.12 MBq*h/MBq. Meanwhile, the gallbladder contents, large intestines, testes, and thymus show relatively shorter residence time ranging from ∼0.00011 MBq*h/MBq to ∼ 0.00051 MBq*h/MBq.

**Table 3.** Residence times of [^11^C]3MeO4AP for measured organs and remainder of body.

| Organ | Residence Time (MBq*h/MBq) |  |
| --- | --- | --- |
|  | Male | Female |
| Adrenals | $4.73 \times 10^{-04}$ | $6.09 \times 10^{-04}$ |
| Brain | $2.02 \times 10^{-02}$ | $1.92 \times 10^{-02}$ |
| Breasts | $1.33 \times 10^{-03}$ | $2.01 \times 10^{-03}$ |
| Gallbladder Contents | $2.29 \times 10^{-04}$ | $2.26 \times 10^{-04}$ |
| LLI | $6.84 \times 10^{-04}$ | $3.11 \times 10^{-04}$ |
| Small Intestine | $6.61 \times 10^{-03}$ | $8.17 \times 10^{-03}$ |
| Stomach | $6.90 \times 10^{-04}$ | $6.18 \times 10^{-04}$ |
| ULI | $6.84 \times 10^{-04}$ | $3.11 \times 10^{-04}$ |
| Heart Contents | $4.22 \times 10^{-02}$ | $3.37 \times 10^{-02}$ |
| Heart Wall | $3.53 \times 10^{-03}$ | $2.94 \times 10^{-03}$ |
| Kidneys | $4.17 \times 10^{-02}$ | $4.04 \times 10^{-02}$ |
| Liver | $5.36 \times 10^{-02}$ | $5.25 \times 10^{-02}$ |
| Lungs | $8.58 \times 10^{-03}$ | $4.40 \times 10^{-03}$ |
| Muscle | $1.48 \times 10^{-01}$ | $1.06 \times 10^{-01}$ |
| Ovaries | NA | $8.69 \times 10^{-05}$ |
| Pancreas | $1.39 \times 10^{-03}$ | $1.81 \times 10^{-03}$ |
| Red Marrow | $1.49 \times 10^{-03}$ | $4.65 \times 10^{-03}$ |
| Cortical Bone | $7.13 \times 10^{-03}$ | $1.30 \times 10^{-02}$ |
| Trabecular Bone | $1.87 \times 10^{-03}$ | $1.89 \times 10^{-03}$ |
| Spleen | $4.20 \times 10^{-03}$ | $3.24 \times 10^{-03}$ |
| Testes | $3.59 \times 10^{-04}$ | NA |
| Thymus | $9.02 \times 10^{-05}$ | $1.26 \times 10^{-04}$ |
| Thyroid | $4.69 \times 10^{-04}$ | $5.14 \times 10^{-04}$ |
| Urinary Bladder Contents | $8.91 \times 10^{-04}$ | $5.19 \times 10^{-03}$ |
| Uterus/Uterine Wall | NA | $1.61 \times 10^{-03}$ |
| Remainder | $8.64 \times 10^{-02}$ | $1.16 \times 10^{-01}$ |

From the residence times, organ dosimetry data was calculated (Table 4). The kidneys (maximum dose: 39.25±1.48 μGy/MBq), heart wall, liver, and adrenals show the highest dose compared with other organs. Meanwhile, the kidneys (maximum contribution: 0.982 ± 0.038 μSv/MBq), liver, lungs, and testes contribute the most to effective dose, which was 4.27 ± 0.57 μSv/MBq based on OLINDA calculation. The male animal effective dose was consistent with the female data. The effective dose of carbon-11 tracer [^11^C]3MeO4AP is notably lower than the fluorine-18 4AP analogue [^18^F]3F4AP (measured in human: 12.2 ± 2.2 μSv/MBq [20]; measured in NHPs: 21.6 ± 0.6 μSv/MBq [12]) but comparable to the effective dose of other carbon-11 tracers measured in humans (e.g. [^11^C]choline: 4.4 μSv/MBq [21], [^11^C]glucose: 4.3 μSv/MBq [22]).

**Table 4.** Organ radiation dosimetry calculations of [^11^C]3MeO4AP using OLINDA software.

| Target Organ | Organ dose<br>($\mu\text{Gy}/\text{MBq}$) | EDE Cont.<br>($\mu\text{Sv}/\text{MBq}$) | ED Cont.<br>($\mu\text{Sv}/\text{MBq}$) |
| --- | --- | --- | --- |
| Adrenals | 11.36±2.74 | 0.68±0.16 | 0.028±0.006 |
| Brain | 5.10±0.38 | 0 | 0.012±0.001 |
| Breasts | 2.18±0.41 | 0.32±0.06 | 0.109±0.020 |
| Gallbladder Wall | 3.74±0.42 | 0 | 0 |
| LLI Wall | 2.12±0.16 | 0 | 0.254±0.019 |
| Small Intestine | 4.22±0.90 | 0 | 0.010±0.002 |
| Stomach Wall | 2.52±0.33 | 0 | 0.302±0.041 |
| ULI Wall | 2.46±0.29 | 0 | 0.006±0.001 |
| Heart Wall | 17.15±0.63 | 1.02±0.03 | 0 |
| Kidneys | 39.25±1.48 | 2.35±0.09 | 0.982±0.038 |
| Liver | 11.55±2.05 | 0.69±0.12 | 0.578±0.101 |
| Lungs | 3.46±0.32 | 0.41±0.03 | 0.415±0.039 |
| Muscle | 2.53±0.29 | 0 | 0.006±0.006 |
| Ovaries | 2.57±1.36 | 0.44±0.62 | 0.354±0.500 |
| Pancreas | 7.10±1.41 | 0.24±0.34 | 0.017±0.003 |
| Red Marrow | 2.09±0.44 | 0.25±0.05 | 0.251±0.053 |
| Osteogenic Cells | 3.18±1.18 | 0.09±0.03 | 0.031±0.011 |
| Skin | 1.12±0.22 | 0 | 0.011±0.002 |
| Spleen | 7.90±0.04 | 0.23±0.33 | 0.019±0.001 |
| Testes | 2.99 | 0.74 | 0.597 |
| Thymus | 3.18±0.36 | 0 | 0.007±0.001 |
| Thyroid | 7.27±1.26 | 0.21±0.03 | 0.364±0.063 |
| Urinary Bladder Wall | 4.05±3.09 | 0 | 0.202±0.154 |
| Uterus | 4.17±3.66 | 0 | 0.010±0.009 |
| Total Body | 2.73±0.37 | 0 | 0 |
| Effective Dose Equivalent ($\mu\text{Sv}/\text{MBq}$) | | 7.36±0.60 (male: 6.94; female: 7.79) | |
| Effective Dose ($\mu\text{Sv}/\text{MBq}$) | | 4.27±0.57 (male: 3.87; female: 4.68) | |

### Safety assessment

No changes in vital signs (including temperature, blood pressure and oxygen saturation, heart rate, respiratory rate, and exhaled CO_2_) were observed during the 150 min of PET imaging acquisition. Routine observation in housing during subsequent days revealed no indications of delayed adverse reaction.

## DISCUSSION

The radiosynthesis of [^11^C]3MeO4AP was achieved using the GE TRACERlabT FX C Pro synthesizer. The previously reported semi-automated radiosynthesis method required the manual addition of reagent into the reaction vessel and manually loading the reaction mixture into the prep-HPLC. The operator was exposed to low amounts of radiation during this process and such manual operation does not fulfill the GMP requirement. The newly developed fully automated radiosynthesis process achieved fully remote-controlled synthesis and fulfills cGMP. The final dose passed all the QC tests except the sterility tests which were not performed since radiosynthesis was carried out in non-sterile environment. The total synthesis and purification time of fully automated method is *ca*. 5 minutes shorter than the semi-automated method. Even though, the non-decay corrected radiochemical yield of the automated method is 3% lower, the yield is sufficient to produce doses for human use.

The biodistribution study shows widespread distribution and fast clearance from most organs. The high SUVs of kidneys and urinary bladder confirms that the [^11^C]3MeO4AP undergoes primarily renal clearance and is eventually eliminated in the urine. The whole body biodistribution of [^11^C]3MeO4AP is similar to that of [^18^F]3F4AP, except that there is lower uptake in the stomach. The slower kinetics in the brain compared to other organs are consistent with previous reports and indicate a high level of specific binding in the brain compared to other organs. There were no differences between male and female data.

[^11^C]3MeO4AP shows low radiation dosimetry based on OLINDA calculation. The effective dose is ∼35 % of [^18^F]3F4AP (12.2 ± 2.2 μSv/MBq [20], human data) and similar to [^11^C]choline (4.4 μSv/MBq [21], human data), and [^11^C]glucose (4.3 μSv/MBq [22], human data). A patient receiving ∼ 400 MBq injection dose of [^11^C]3MeO4AP will receive ∼ 1.7 mSv of radiation dose from the PET tracer, which is ∼ 1.7 fold of the annual radiation dose limit of individual members of the public (56 FR 23398, May 21, 1991).

## ACKNOWLEDGEMETNS

We thank David Lee and Hamid Sabet at the MGH Gordon PET cyclotron facility for producing carbon-11. We thank the veterinary staff (Helen Deng) for assistance with animal handling.

We thank Marina Macdonald-Soccorso for running the PET-CT scanner.

## Funding

This study was supported by NIH/NIBIB (MDN and GEF P41EB022544), NIH/NINDS (PB, R01NS114066), NIH/NIBIB (YPZ, 1K99EB033407).

## AUTHOR CONTRIBUTIONS

YPZ contributed to the study design, developed the automated radiosynthesis protocols, synthesized the radiotracers and analyzed the PET imaging and blood data; MQW contributed to the study design, processed and analyzed the dosimetry data; assisted with the PET imaging of NHPs; MD and NG conducted the PET imaging of NHPs; SHM processed the blood samples; GEF contributed to the data interpretation; MDN contributed to the study design, performed NHPs PET scans and supervised the dosimetry study; PB contributed to the study design and supervised the entire project. YPZ and PB wrote the manuscript and all authors reviewed and approved it.

## DISCLOSURES

PB has a financial interest in the Fuzionaire Diagnostics and the University of Chicago. PB is a named inventor of patents related to [^18^F]3F4AP owned by the University of Chicago and licensed to the Fuzionaire Diagnostics. PB’s interests were reviewed and are managed by the MGH and Mass General Brigham in accordance with their conflict of interest policies.

The other authors declare no competing interests.

